# Age-Stage, Two-Sex Life Table of the *Menochilus sexmaculatus* (Coccinellidae: Coleoptera) Feeding on Different Aphid Species

**DOI:** 10.1101/2020.01.15.907576

**Authors:** Khalid Abbas, Muhammad Shah Zaib, Muhammad Zakria, Umm-e -Hani, Syed Muhammad Zaka, Noor-ul -Ain

## Abstract

*Ladybird beetle, Menochilus sexmaculatus (*Fabricius) (*Coleoptera: Coccinellidae*), is biological control agent that predate the different aphid species. Both adults and larval stage of *M. sexmaculatus* feed on aphid species. In this experiment Life table and predation data were collected for *M. sexmaculatus* feed on four different aphid species *Lipaphis erysimi*, *Myzus persicae*, *Aphis nerii* and *Diuraphis noxia*. This experiment was conducted under laboratory conditions at 25±2°C, 60±5% RH and L14: D10 h. Different numbers of aphid were provided as a pray in petri dish. The pre-adult development duration of *M. sexmaculatus* was maximum when fed on *M. persicae* (12.18 d) and minimum on *D. noxia* (10.64 d). Similarly, male and female duration was maximum on *M. persicae* (26.7 d), minimum on *L. erysimi* (23.67 d) in male and in female maximum on *D. noxia* (28.00 d), minimum on *A. nerii* (24.33 d). Net reproductive rate (R_o_) range from 117.9 on *L. erysimi* to 99.55 on *M. persicae* and intrinsic rate of increase (r) range was 0.21197 d^−1^ on *A. nerii* to 0.021559 d^−1^ on *D. noxia*. The finite rate of increase (λ) range was 1.240592 d^−1^ on *D. noxia* to 1.204918 d^−1^ on *M. persicae*, the mean of generation (T) range was 24.68 d^−1^ on *M. persicae* to 22.476 d^−1^ on *A. nerii*, similarly, the gross reproductive rate (GRR) range was 172.2 d^−1^ on *D. noxia* to 115.02 d^−1^ on *M. persicae* and Fecundity (F) eggs per female range was 316.8 on *D. noxia* to 199.1 on *M. persicae*. In present Study, age-stage two-sex life table gives complete understanding of predator biological aspects against different aphid species. This study will help us to improve mass rearing and use of *M. sexmaculatus* in biological control of aphids.

## Introduction

Aphids (Hemiptera: Aphididae) are important insect pests of various cultivated plants (1). Suck cell sap of plants and act as vectors of various virus induced diseases (2). They have abilities to quickly build their population and their honeydew secretions results into a medium of sooty mold growth. They can change host metabolism by disturbing their host hormonal balance. Aphids attack may kill the plant at their early growth stages and reduce yield of crops at later stages (3). Oleander aphid (*Aphis nerii*), green peach aphid (*Myzus persicae*), Russian wheat aphid (*Diuraphis noxia*) and mustard aphid (*Lipaphis erysimi*) are among important pests of cultivated and ornamental plants (4). The *L. erysimi*, most importantly damages *Brssicace* plants typically mustard, rape, cabbage, cauliflower, broccoli and radish worldwide (5). The *M. persicae*, is a cosmopolitan pest, feeds on more than 50 plant families (5), including agro-industrial crops and horticultural crops (6).

The *D. noxia*, attacks on cereal crops worldwide with high host range of more than 140 species of Poaceae plants (7). The *D. noxia*, inject toxin into plants while feeding which causes failure to unrolling and white streaking of plant leaves. Yield loss had been estimated up to 80 to 100% under heavy attack of *D. noxia*, in wheat crop (8). The *A. nerii*, feeds on plants of Apocynaceae and Asclepiadaceae families (9) and also had been reported on wheat and Brassica in Pakistan (10). The *A. nerii*, is an obligate parthenogen, and a sequester of toxic chemicals (cardenolides) which act as defensive mechanism against its natural enemies (11). Indeed, unjudicious pesticides use increased ability of pests to survive against pesticides and residues level in crops final produce ((12) (13) and these factors urge to use alternative methods (e.g. biological control) to reduce aphid populations which are environmental friendly and risk free for human health.

Natural enemies (predators, parasitoids and entomopathogens) used to control aphids population in biological control (14). Natural enemies are the basic components of insect pest supervision. Practically 90% of natural pests are controlled by natural enemies (15). Ladybirds are potent predators of various small herbivorous insects such as aphids (16). The Ladybird beetle, *Menochilus sexmaculatus* (Fab.), is distributed in Pakistan, India and other south Asian countries (17). The adults of *M. sexmaculatus* are yellow bright in color and having black zigzag lines. Some preys are toxic to predators because they feed on toxic plant and ultimately affects food quality for predators (18). Few studies have been done on biological aspect of *M. sexmaculatus* against different aphid species. However, there is a need for detail study of survival and reproduction of M. sexmaculatus on aphid species to evaluate suitable prey and alternate prey species. It is important to know demographic aspects including stage differentiation and predation rate of predators for mass rearing of predators and true implication into biological control of pests (19). Therefore, life table was studied to know the development and reproduction of predators against pests. However, age-stage two-sex life table provides more detail of biological aspects including stage differentiation than traditional life tables (19). Therefore, present study used age-stage two-sex life table for complete understanding of *M. sexmaculatus* biological aspects against different aphid species. This study will help us to improve mass rearing and use of *M. sexmaculatus* in biological control of aphids.

## Material and Methods

### Rearing of Aphids

Four aphid species (*A. nerii M. persicae*, *D. noxia* and *L. erysimi*) were collected from their hosts from agricultural fields (latitude 30°15’29.9"N, longitude 71°30’54.6"E) of Faculty of Agricultural Sciences and Technology, Bahauddin Zakariya University, Multan Pakistan and were reared on their respective host plants. Aphids were reared in plastic cages (51 **×** 45 cm) along with their respective hosts under laboratory condition (25 ± 2°C and 70 ± 5% RH with photoperiod of 14L:10D h) (20). This laboratory reared aphids were used for the biological studies of *M. sexmaculatus*.

### Collection and rearing of *M. sexmaculatus*

The larvae of *M. sexmaculatus* were collected from *Calotropis procera* located at head Muhammad wala fields of Multan (latitude: 30°11′54.97N, longitude: 71°28′7.33E), Punjab, Pakistan in start of February 2019. Larvae were collected in early morning in plastic jars (25 × 15.5 cm) with the help of camel hairbrush and transferred to aphid culture in laboratory. The culture was maintained in an incubator (25±1°C and 60±2% R.H.) with photoperiod 14L:10D h) (21). The collected larvae were transferred to plastic jars (15 × 11 cm). The mouth of cages was covered with the muslin cloth and knotted with the elastic band. Different aphid species i.e. *A. nerii*, *M. persicae*, *D. noxia* and *L. erysimi* were supplied as a food to larvae. Emerging adults were reared in plastic boxes (14 × 8 × 10 cm) with surfeit different aphid species. Corrugated filter papers were used as an oviposition substrate of beetles in rearing boxes. Collected eggs from these adult females were placed in 10-cm petri dishes containing moist filter paper at the bottom to get larvae. Mature and immature stages of *M. sexmaculatus* were provided with aphids as their food (22).

### Life table Studies

Fifty healthy eggs of *M. sexmaculatus* were taken from the general papulation of their respective hosts and kept separately in single petri dishes (6cm diameter). Egg development period was recorded after 6-h interval. After egg hatching 1^st^ instar larvae of *M. sexmaculatus* were feed on aphid species and similarly all instar of *M. sexmaculatus* were feed on aphid species. Specified number of aphids were provided, and data of consumed aphids were recorded on daily basis (23). After 4^th^ instar larvae convert into the pupal stage and then into the adult stage. Duration of All stages (larvae pupae and adult) were recorded 12-h interval (20, 21, 24). Adult male and female were paired in plastic jars (9 × 6 cm) for mating, egg laying and to check the male and female longevity, reproductive behavior and female oviposition. Similarly, male and female were kept separately to check the predation rate and observe the fecundity and survival rate of both sexes were recorded after 24-h until death (24, 25). The software TWOSEX-MS Chart (26) was use to check the egg to adult development duration, fecundity, adult preoviposition period, oviposition period, post oviposition period and age two sex life cycle (27, 28). Age-specific survival rates were find according to (27) life expectancy according to (19) and papulation growth on different aphid species.

### Statistical analysis

Development duration and population parameters were calculated using TWOSEX-MS Chart, to minimize variation in the results. The bootstrap technique (29) with 100,000 replications was used to calculate the mean and SE of the population (30). The TIMING-MS Chart program (31) based on age-stage two sex life table for data of *M. sexmaculatus*. The raw data were used to calculate the age-stage–specific survival rate (*s*_*xj*_, where *x* = age in days and *j* = stage), age-stage specific fecundity (*f*_*xj*_), age-specific survival rate (*l*_*x*_), age-specific fecundity (*m*_*x*_), age-specific net maternity (*l*_*x*_*m*_*x*_), age-stage life expectancy(*e*_*xj*_), age-stage reproductive value (*v*_*xj*_), and life table parameters (32) (*R*_0_,net reproductive rate; *r*, intrinsic rate of increase; *λ*, finite rate of increase; and *T*, the mean generation). In the age-stage, two-sex life table, the age-specific survival rate *l*_*x*_, *m*_*x*_ and *R*_0_ was calculated as (1 and 2):

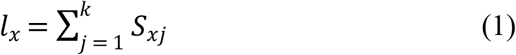

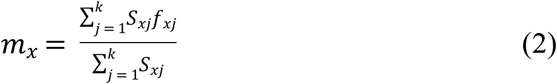

Where *k* is the number of stages. The net reproductive rate *R*_0_ is the mean number of offspring laid by individual during its entire life span. It was calculated by following equation (3):

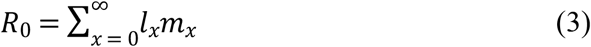

The intrinsic rate of increase (*r*) was estimated using the iterative bisection method and corrected with the Euler–Lotka equation (4) with the age indexed from 0 (33):

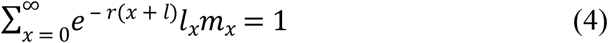

The finite rate (*λ*) was calculated as (5):

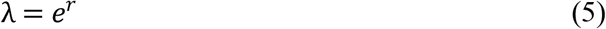

The mean generation time is defined as the length of time that a population needs to increase to *R*_0_-fold of its population size at the stable age-stage distribution, and is calculated as (6):

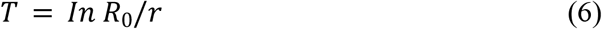

The life expectancy (*e*_*xj*_) is the length of time that an individual of age *x* and stage *j* is expected to live and it is calculated equation (7) according to as (19).

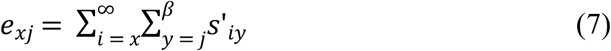

The comparison between different aphid species were done by using completely randomized design and means were compared by using LSD test (P=0.05). This analysis was done by using statistical package SAS (34).

## Results

When different aphid’s species were given to immature stages of beetle, significant (P=0.0032, F=0.13 and DF=3) different response on survival was recorded (Table 1) i.e. highest survival (89.1) was recorded when *L. erysimi* was given as a diet. While *A. nerii*, *M. persicae* and *D. noxia* gave similar result (85, 85 and 84.1, respectively) for immature survival.

**Table 1.** Development period parameters (mean ± SE) of *M. sexmaculatus*

| Aphid species | *n* | *Aphis nerii* | *Myzus persicae* | *Diuraphis noxia* | *Lipaphis erysimi* | *pvalue* | *f.value* | *df* |
| --- | --- | --- | --- | --- | --- | --- | --- | --- |
| Immature survival (%) | 20 | 85.0±6.4b | 85±7.3b | 84.1±6.7b | 89.1±5.0a | 0.0032 | 0.13 | 3 |
| Adult emergence (%) | 20 | 90.0±6.9b | 90±6.9b | 95±9a | 90±6.9b | 0.0054 | 0.15 | 3 |
| Developmental rate (d<sup>-1</sup>) | 20 | 00.04c | 00.04c | 00.04b | 00.06a | <0.0001 | 2.15 | 3 |
| Male longevity (d) | 20 | 24.30±4.7b | 26.7±3.53a | 26.33±2.2a | 23.67±3.7c | <0.0001 | 0.19 | 3 |
| Female longevity (d) | 20 | 24.30±5.9b | 25.7±5.5b | 28±3.1a | 27.33±0.9b | <0.0001 | 0.16 | 3 |
| Pre-oviposition period (d) | 10 | 07.0±2.5a | 6.33±0.9b | 5.33±1.3b | 5.67±1.5b | 0.0211 | 0.18 | 3 |
| Oviposition period (d) | 10 | 15.70±2.3b | 18.33±1.9a | 14.70±1.3b | 10.33±6.9c | <0.0001 | 0.87 | 3 |
| Post-oviposition period (d) | 10 | 11.00±3.2a | 11.33±3.8a | 7.33±1.7c | 9.70±4.3b | <0.0001 | 0.72 | 3 |

n= number of replication, Mean value; SE, standard error; df, degree of freedom; *F*, value by statistical package SAS; *P*, statistical significance level 0.05. Mean followed by different letters in the same row are significantly different by statistical package SAS using of difference.

The adult emergence was recorded, significant (P=0.0054, F=0.15 and Df=3) when their immature stages fed on different aphid species (Table 1) i.e. maximum adult emergence (95.00 %) was recorded when fed on *D. noxia* but when fed on *A. nerii*, *M. persicae* and *L. erysimi* (90, 90 and 90 %, respectively) the adult emergence recorded was similar.

When different species of aphid were offered to immature stages of beetle, significant (P<0.0001, F=2.15 and Df=3) difference was recorded in developmental rate (Table 1) i.e. maximum developmental rate was recorded (0.057 d^−1^) on *L. erysimi* followed by *D. noxia* (0.042 d^−1^). While, similar result of developmental rate (0.038 and 0.035 d^−1^, respectively) was observed when immatures were fed on *A. nerii* and *M. persicae*.

The significant difference in adult longevity of both males and females was observed when different aphid species were provided as a diet (Table 1). The significantly (P<0.0001, F=0.19 and Df=3) maximum male longevity was observed on *M. persicae* and *D. noxia* (26.7 and 26.33 d, respectively), followed by A*. nerii* (24.33 d). While minimum male longevity was observed when *L. erysimi* was given as a diet (23.67 d). In case of female, significant response was recorded (P<0.0001, F=0.19 and Df=3), maximum longevity was recorded on *D. noxia* (28.00 d) followed by *L. erysimi*, *M. persicae* and *A*. *nerii* (27.33, 25.7 and 24.33 d, receptively).

When beetle was provided different aphid species as a diet, significant (P=0.0211, F=0.18 and Df=3) difference in the pre-oviposition period was recorded (Table 1) i.e. maximum pre-oviposition period of beetle was recorded when fed on A*. nerii* (7.00 d). While, when *M. persicae*, *L. erysimi* and *D. noxia* (6.33, 5.67 and 5.33 d, respectively) were provided as a diet to beetle showed same result.

The oviposition period of beetle, significant (P<0.0001, F=0.87 and Df=3) difference was recorded when they fed on different aphid species (Table 1), i.e. highest oviposition period was recorded (18.33 d) when fed on *M. persicae*. The oviposition period of beetle was recorded (15.70 and 14.70 d, respectively) similar when they fed on *A. nerii* and *D. noxia*, respectively, and followed by *L. erysimi* (10.33 d).

The post-oviposition period of beetle, significant (P=<0.0001, F=0.72 and Df=3) difference was observed when they were provided four different aphid species (Table 1) i.e. maximum post-oviposition was recorded (11.33 and 11.00 d, respectively) on *M. persicae* and *A. nerii* and followed by *L. erysimi* (9.7 d). While minimum post-oviposition period was observed (7.33 d) on *D. noxia*.

When different aphid species were provided to *M. sexmaculatus* the significant (P=0.0146, F=5.15 and Df=3) difference in incubation period was recorded (Table 2) i.e. maximum incubation period was noted on *M. persicae* (2.53 d), followed by *A. nerii*, *L. erysimi* and *D. noxia* (2.23, 2.10 and 2.04 d, respectively).

**Table 2.**
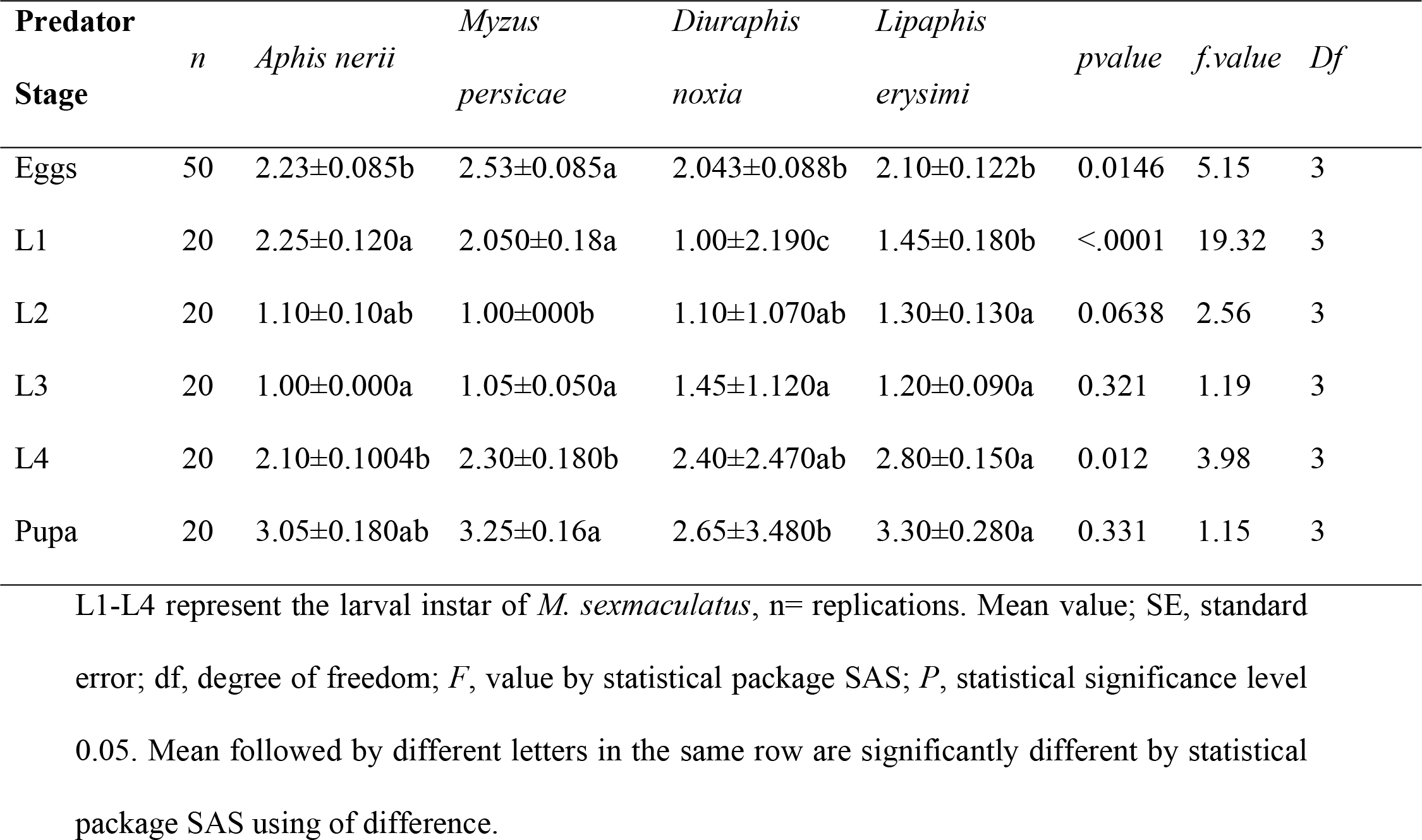
Immature developmental time (mean ± SE) of *M. sexmaculatus*

Different species of aphid were given to different larval stages of *M. sexmaculatus*, the significant difference in developmental time was recorded (Table 2). The significantly (P<0.0001, F=19.32 and Df=3) highest first instar (L1) developmental time was recorded on *A. nerii* and *M. persicae* (2.25 and 2.05 d, respectively), followed by *L. erysimi* (1.45 d). While shortest developmental time was recorded on *D. noxia* (1.00 d). The maximum significant (P=0.0638, F=2.56 and Df=3) developmental time of second instar (L2) was recorded on *L. erysimi* (1.30 d) followed by *A. nerii* and *D. noxia* (1.10 and 1.10 d, respectively). Whereas minimum developmental time was recorded on *M. persicae* (1.00 d). The developmental time of third instar (L3) observed was non-significant (P=0.321, F=1.19 and Df=3) on all four aphid species. The highest significant (P=0.012, F=3.98 and Df=3) developmental time of fourth instar (L4) was noted on *L. erysimi* (2.80 d) followed by *D. noxia* (2.40 d). The lowest developmental time was recorded on *M. persicae* and *A. nerii* (2.30 and 2.10 d, respectively).

The developmental time of pupae on all four aphid species was recorded non-significant (P=0.331, F=1.15 and Df=3) (Table 2).

When different species of aphid were provided the significant difference in intrinsic rate of increase (r) was recorded (Table 3) i.e. maximum intrinsic rate of increase (0.21197 d^−1^) when fed on *A. nerii* and followed by *L. erysimi* and *M. persicae* (0.198695 and 0.186412 d^−1^, respectively). While minimum intrinsic rate of increase (r) was recorded (0.021559 d^−1^) when fed on *D. noxia*.

**Table 3.** Life table parameters mean of *M. sexmaculatus*

| Aphid species | *Aphis nerii* | *Myzus persicae* | *Diuraphis noxia* | *Lipaphis erysimi* |
| --- | --- | --- | --- | --- |
| r (d<sup>-1</sup>) | 0.21197 | 0.186412 | 0.021559 | 0.198695 |
| λ (d<sup>-1</sup>) | 1.236111 | 1.204918 | 1.240592 | 1.21981 |
| R<sub>0</sub>(Offspring individual<sup>-1</sup>) | 117.25 | 99.55 | 158.4 | 117.9 |
| T (d) | 22.476 | 24.68 | 23.494 | 24.006 |
| GRR | 131.92 | 115.02 | 172.2 | 125.67 |
| F | 260.56 | 199.1 | 316.8 | 235.8 |

$r$ = intrinsic rate of increase, $\lambda$ = finite rate of increase, $R_0$ = net reproductive rate $T$ = the mean of generation, $GRR$ = the gross reproductive rate and $F$ = Fecundity (eggs per female).

The significant difference in finite rate of increase (*λ*) was recorded when different aphid species were given to *M. sexmaculatus* (Table 3) i.e. maximum finite rate of increase (*λ*) was reported (1.240592 d^−1^) when fed on *D. noxia* followed by (1.236111 and 1.21981 d^−1^, respectively) when fed on *A. nerii* and *L. erysimi*, respectively. Minimum finite rate of increase was recorded when fed on *M. persicae* (1.204918 d^−1^).

When different aphid species were given to *M. sexmaculatus* the significant difference in net reproductive rate (R_o_) was recorded (Table 3) i.e. maximum net reproductive rate (R_o_) was recorded (158.4 d^−1^) when fed on *D. noxia* followed by *L. erysimi* and *A. nerii* (117.9 and 117.25 d^−1^, respectively), whereas minimum net reproductive rate (*R*_*o*_) recorded when fed on *M. persicae* (99.55 d^−1^).

The significant difference in mean of generation (*T*) was recorded when different aphid species were provided to *M. sexmaculatus* (Table 3) i.e. maximum mean of generation (*T*) was reported (24.68 d^−1^) when fed on *M. persicae* fallowed by (24.006, 23.494 d^−1^, respectively) *L. erysimi*, *D. noxia* respectively. While minimum mean of generation (*T*) was recorded (22.476 d^−1^) when fed on *A. nerii*.

The significant difference in gross reproductive rate (*GRR*) of *M. sexmaculatus* was observed when different aphid species were provided (Table 3) i.e. maximum gross reproductive rate (*GRR*) was recorded (172.2 d^−1^) when fed on *D. noxia* fallowed by (131.92 and 125.67 d^−1^, respectively) *A*. *nerii* and *L. erysimi*, respectively. While minimum gross reproductive rate (*GRR*) was reported (115.02 d^−1^) when fed on *M. persicae*.

When different aphid species were given to *M. sexmaculatus* the significant difference was observed in fecundity (F) i.e. maximum fecundity (F) was recorded (316.8) when fed on *D. noxia* followed by (260.56 and 235.8, respectively) *A. nerii* and *L. erysimi* respectively. While minimum fecundity (F) was recorded (199.1) when *M. persicae* was given (Table. 3).

Age-stage-specific survival rate (*s*_*xj*_) curves (Fig 1) show that stage survival curves are overlapping with each other due to difference in developmental duration. *M. sexmaculatus* when feed on *M. persicae* show maximum survival to adult stage than *D. noxia*, *L. erysimi* and *A. nerii*. Whereas adult survival of *M. sexmaculatus* was similar in *M. persicae* and *D. noxia*.

**Fig 1.**
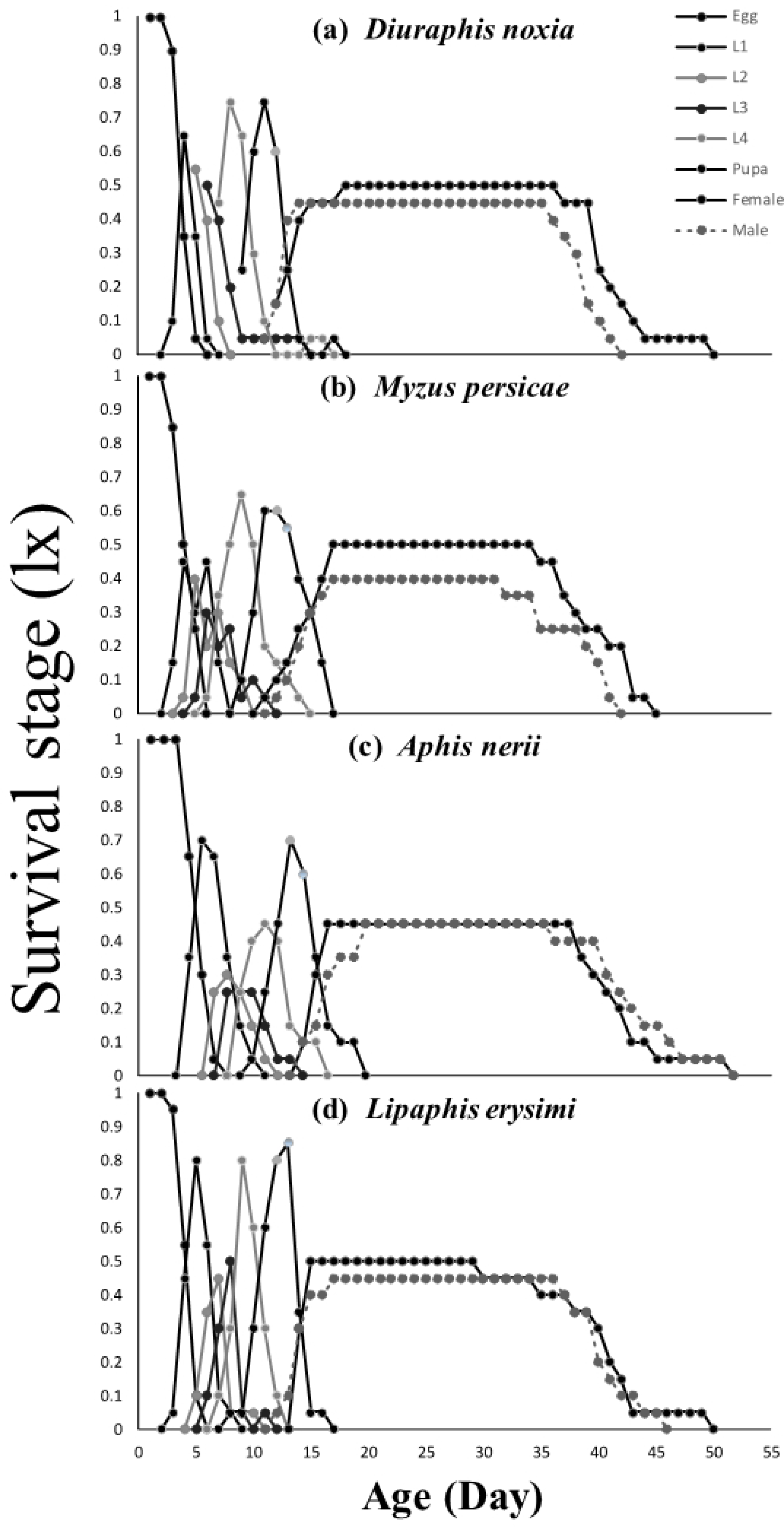
Age-stage–specific survival rate (*s*_*xj*_) of *M. sexmaculatus* fed on four aphid species.

*M. sexmaculatus* evinced similar but maximum survival rate both on *M. persicae* and *D. noxia* according to age specific survival rate (Fig 2). The age-stage-specific female fecundity (*f*_*x7*_) and age-specific fecundity (*m*_*x*_) shows that beetle maximum oviposition was 29.4 eggs at age of 23 days (fig. 2) and 15.5 eggs, respectively (Fig 2). The values of (*f*_*x7*_) and (*m*_*x*_) of beetle were minimum on turnip aphids. The age-specific net maternity (*l*_*x*_*m*_*x*_) shows that highest age-specific net maternity (*l*_*x*_*m*_*x*_) was recorded on *D. noxia* followed by *L. erysimi* and *A. nerii*. Whereas minimum was recorded on *M. persicae*.

**Fig 2.**
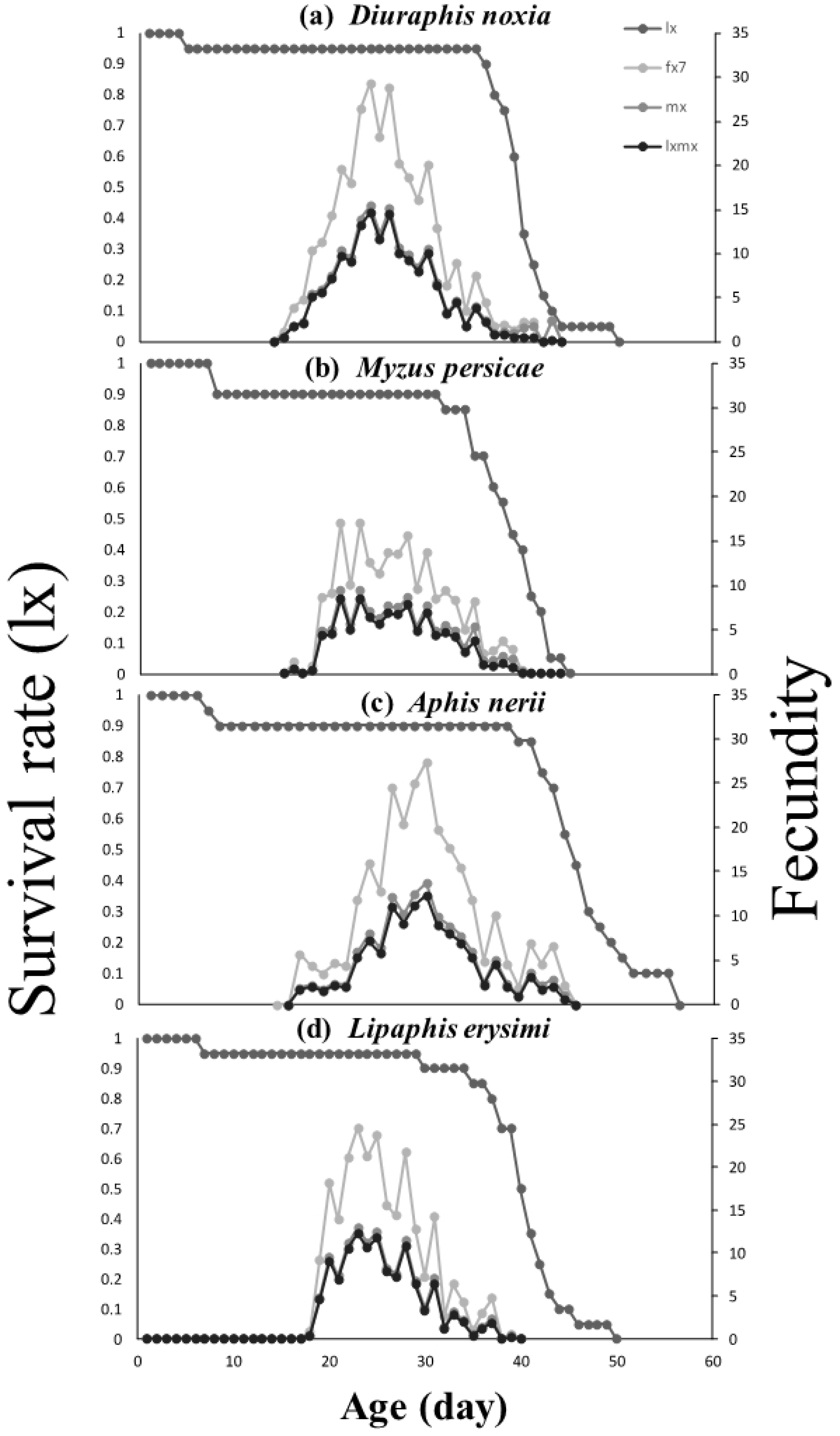
Age-specific survival rate (*l*_*x*_), age-stage–specific fecundity (*f*_*xj*_), age-specific fecundity (*m*_*x*_), and age-specific maternity (*l*_*x*_*m*_*x*_) of *M. sexmaculatus* fed on three aphid species.

Age-stage–specific reproductive rates (*v*_*xj*_) shows (Fig 3) that it is highest in case of *D. noxia* (110) at the age of 22 days. The highest reproductive values *A. nerii L. erysimi* and *M. persicae* are 98 at 21days, 96 at 22 days, and 73 at 20 days, respectively.

**Fig 3.**
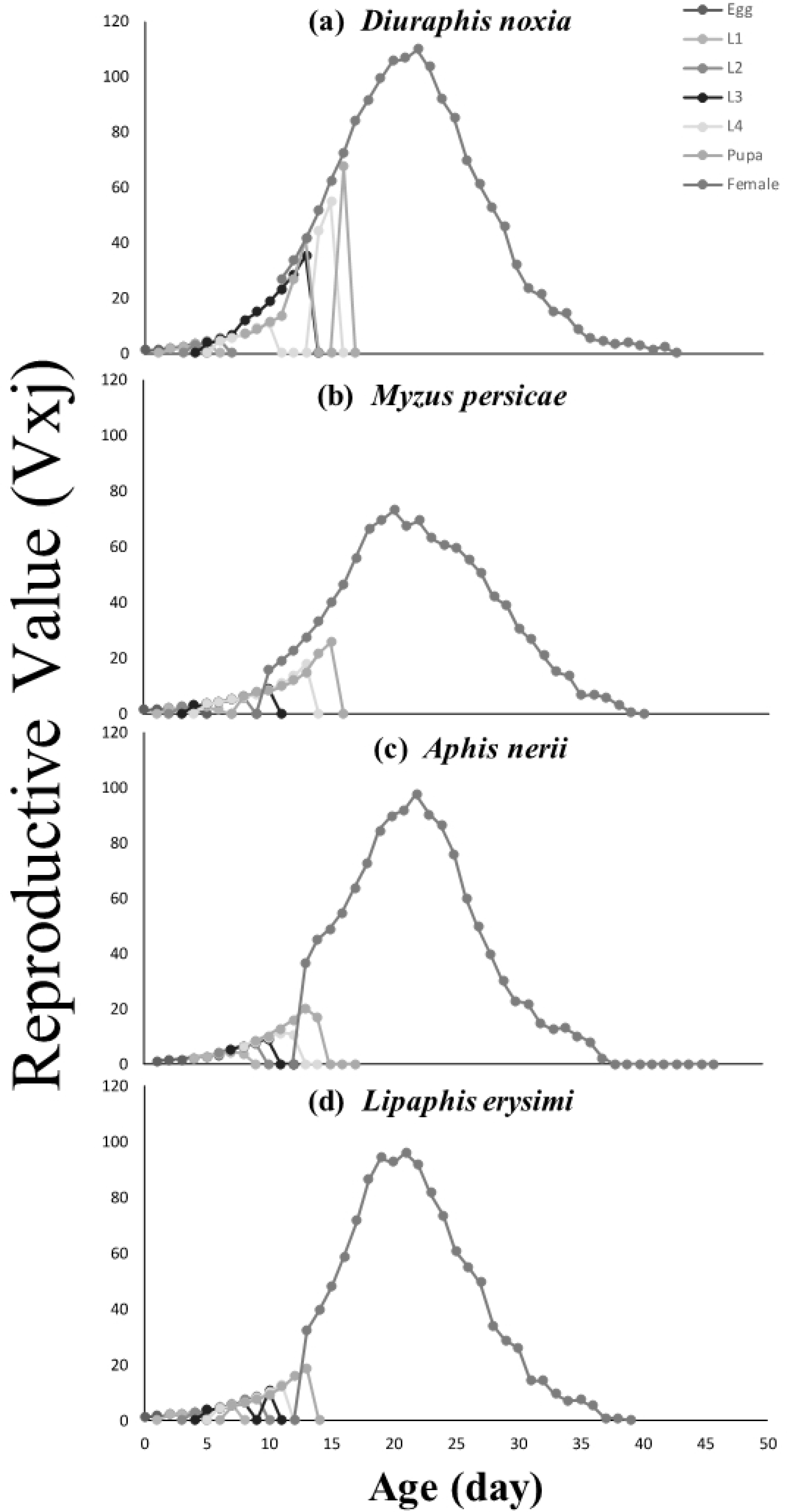
Age-stage–specific reproductive rate (*v*_*xj*_) of *M. sexmaculatus* fed on four aphid species.

Life expectancy curves (*e*_*xj*_) of females are similar in case of *D. noxia* and *L. erysimi* however are larger than *M. persicae* and *A. nerii*. Life expectancy curves presented (Fig 4) the survival of individual age *x* and stage *j*. Freshly hatched eggs of *M. sexmaculatus* estimated to live for 35, 35, 34.5 and 32.5 days on *M. persicae*, *L. erysimi*, *D. noxia* and *A. nerii*, respectively. Usually female life expectancy greater than male life expectancy but in case of *A. nerii* and *L. erysimi* male life expectancy was greater than female life expectancy. Female and male life expectancies were reported 30 and 28 days after age of 12.5 and 10 days on *D. noxia*, respectively, while greater than *M. persicae* (29 and 26 days after age of 11 and 11.5 days, respectively).

**Fig 4.**
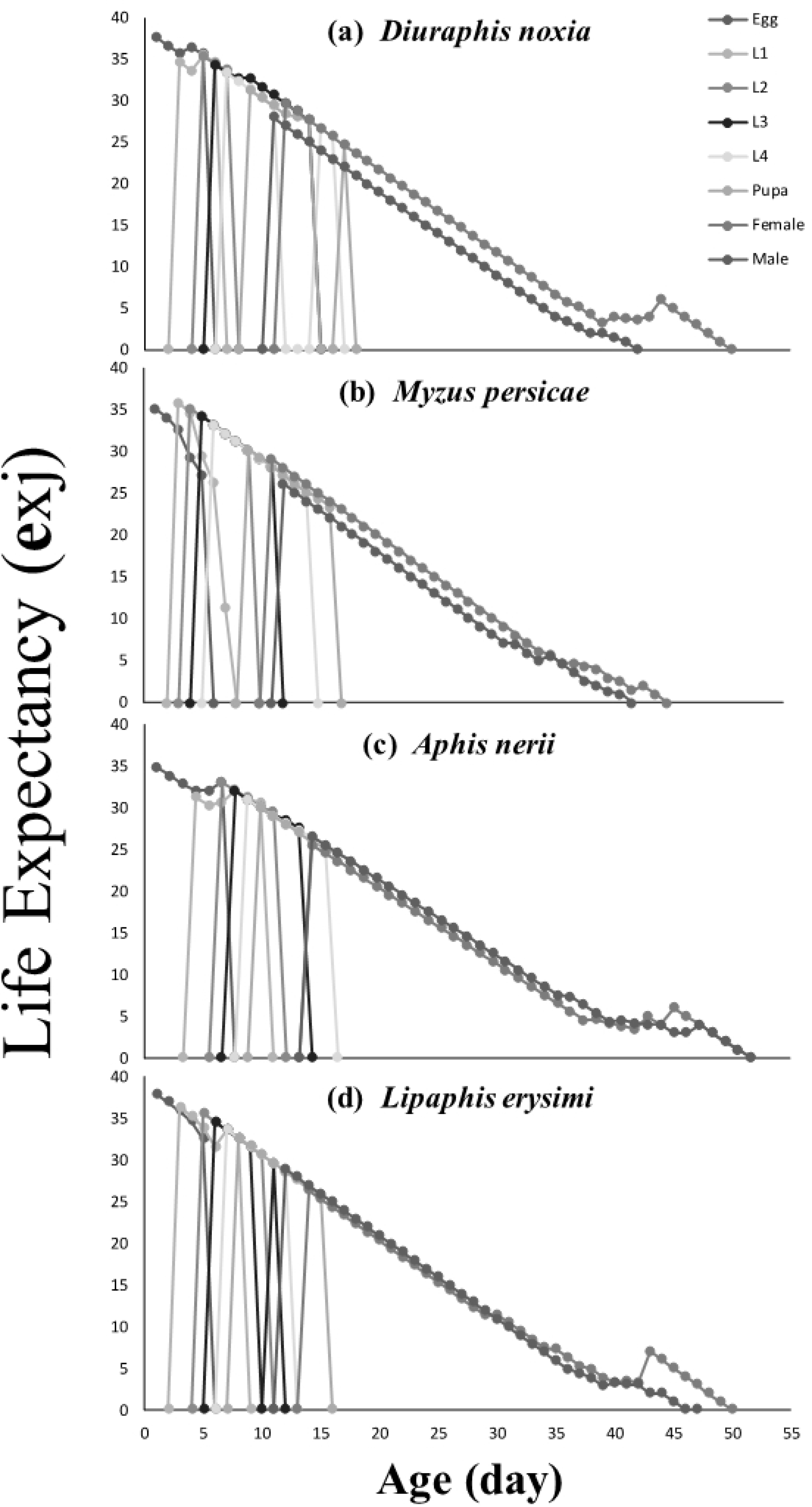
Age-stage–specific life expectancy (*e*_*xj*_) of *M. sexmaculatus* fed on four aphid species.

## Discussion

*M. sexmaculatus* is good predator of aphids and an important biological control agent. The present study was carried out to understand the effect of different aphid species on the development, fecundity and survival rate of *M. sexmaculatus*. The results of present study showed that quality and availability of prey affect the development of *M. sexmaculatus*. These results closely related with work of (35) who reported that the quality and nature of the prey affect the development, fecundity and survival rate of predator. Low quality and insufficient quantity of prey reduce the development of predator, whereas good quality and enough quantity of prey increase the development of predator (36).

Results showed that on comparison between *L. erysimi* and *M. persicae*, the maximum male longevity was recorded on *L. erysimi* while minimum was recorded on *M. persicae*. These results correlate with the study (24, 37) where *C. septempunctata* males exhibited maximum longevity on *L. erysimi* as compared to *M. persicae*. While in case of female, maximum longevity was recorded on *M. persicae* as compared to *L. erysimi*, this contradict with the result of *C. septempunctata* female population which showed maximum longevity on *L. erysimi* then *M. persicae*. This might be due to different species of beetles.

The results of present study revealed that statistically maximum male and female longevity was recorded on *D. noxia* while minimum longevity was recorded on *L. erysimi*. These results contrary with the study conducted on *C. septempunctata* that the adult longevity was maximum on *L. erysimi* than other aphid species (24, 38). In current study, highest fecundity was recorded on *D. noxia*. These results contrary with the study carried out on *C. septempunctata* where the maximum fecundity was reported on *M. persicae* (24, 39). There is a relation among predator longevity and fecundity. The predator has long longevity it does not mean that they have maximum fecundity. Because quality of host affects the longevity and fecundity of predator (39, 40).

The results of current study revealed that maximum age stage specific survival rate (*s*_*xj*_) was recorded on *M. persicae*. These results resembled with the study conducted on *C. septempunctata* that the maximum survival rate was recorded on *M. persicae* (19, 24, 41, 42). In this study maximum developmental rate was observed on *L. erysimi*. These findings closely resembled with the study performed on *C.* septempunctata. Which also showed maximum developmental rate was on *L. erysimi* as compared to other aphid species. The reason was that the quality and quantity of prey affect the developmental rate of both immature and adult stages (43).

The biological parameters of predator are heavily affected by several factors like type of prey. The findings of current study revealed that the maximum *R₀* and *λ* was recorded on *D. noxia*. The maximum *r* was recorded on *A. nerii*. The highest *T* was noted on *M. persicae*. These results contrary with the study conducted on *C. septempunctata* that the maximum *R₀*, *λ* and *r* was recorded on *M. persicae*, whereas maximum *T* was recorded on *L. erysimi* (24, 44, 45). The results of present study revealed that the maximum TPOP was recorded on *A. nerii*. These results contradict with the study performed on *C. septempunctata* that the maximum TPOP of *C. septempunctata* was recorded on *L. erysimi* (21, 24). In laboratory conditions TPOP of *M. sexmaculatus* was recorded minimum by Zhao et al. (25). The reason was that the difference in biotic and abiotic factors are responsible for changes in the findings (44).

In previous studies problems were associated with the traditional life table i.e. consider female population, neglect male population and stage differentiation between individuals and sexes. In present study age stage two sex life table was used to assess the difference between age specific survival rate and age specific fecundity which also consider the male survival curve and stage differentiation between individuals. The difficulties and errors associated with the female age specific life table briefly addressed by (19, 46).

The results of present study showed that oviposition period was maximum when they fed on *M. persicae*. These results contradict with the study conducted on *C. septempunctata* that the maximum oviposition period was recorded on *L. erysimi* (24, 47). The results of present study revealed that the maximum fecundity curve (29.4 eggs) was reported on 23^rd^ day, daily and lifelong fecundity were recorded on *D. noxia* (23.70 and 110 eggs, respectively). These results contradict with the study conducted on *C. septempunctata* that the maximum fecundity curve was reported (36.111 eggs) on 43^rd^ day, daily and lifelong fecundity (39 and 470 eggs, respectively) were reported on *M. persicae* (24, 47, 48). The reason was that the nutritional value and quality of prey species affect the predator fecundity (49, 50). The life expectancy is that an adult is supposed to live at age *x* and stage *j*. The results of this study expressed that the life expectancy was reduced with the age of an adult. These results resembled with the study conducted on *C. septempunctata* that the adult’s life expectancy reduced with the age. Without giving any stress adult’s life expectancy gradually reduced with the age under laboratory conditions (24, 51, 52). The life expectancies of same age individuals can be changed, by the difference in life stages of individuals (19).

The current study was designed to evaluate the population growth in association with the number of individuals instead of *r*. That provides evidence about the growth potential of a population at an even age distribution (53). It was intended that *M. sexmaculatus* reached a stable age stage after 23 days when reared on *D. noxia*. The maximum population was observed on *D. noxia* as compared to other species. It is reflected that *D. noxia* is most suitable host for mass rearing of *M. sexmaculatus* under laboratory conditions.

## Conclusion

It was concluded that the prey specificity and availability affect the life table parameters of *M. sexmaculatus*. The appropriate host for mass rearing of *M. sexmaculatus* is *D. noxia* under laboratory conditions. Moreover, both male and female includes in age-stage two-sex life table. Because age-stage two-sex life table gives brief information about the efficacy and use of *M. sexmaculatus* population in biological control. Future studies should consist on field application and evaluation of *M. sexmaculatus* for the management of aphid.

## Acknowledgment

The authors would like to thank Mr. Yasir Hameed, Muhammad Sarmad and Muhammad Farrukh Hamid for their help during the work. Moreover, we grateful to Department of Entomology, Faculty of Agricultural Sciences & Technology, Bahauddin Zakariya University Multan for providing support and facilities to perform the Research.

